# How do life history traits influence the environment’s effect on population synchrony? Insights from European birds and insects

**DOI:** 10.1101/2023.09.08.556676

**Authors:** Ellen C. Martin, Brage Bremset Hansen, Aline Magdalena Lee, Ivar Herfindal

**Author notes:** Corresponding author: Ellen C. Martin.

## Abstract

Populations closer together in space are more likely to experience shared environmental fluctuations. This correlation in experienced environmental conditions is the main driver of spatial population synchrony, defined as the tendency for geographically separate populations of the same species to exhibit parallel fluctuations in abundance over time. Moran’s theorem states that spatially distinct populations are expected to show the same synchrony in their population dynamics as the synchrony in their environment. However, this is rarely the case in the wild, and the population synchrony of different species inhabiting the same area is rarely similar. These species-specific differences in how the environment synchronizes populations can be due to life history traits that make some species more susceptible to environmental stochasticity, such as reduced mobility or faster pace of life. In this study, we compiled long-term annual abundance datasets on European birds and insects (*Lepidoptera* sp. and *Bombus* sp.) to identify how environmental synchrony (i.e., positively spatially correlated fluctuations in the environment, also called the Moran effect) affects species population synchrony. As expected, the environment synchronized populations of both birds and insects. Populations experiencing correlated fluctuations in precipitation or temperature had higher synchrony in annual population growth rates. Birds were more strongly synchronized by temperature, while precipitation was a stronger driver of synchrony in insects. In birds, species with short generation times had a stronger synchronizing effect of the environment compared to species with long generation times. Moreover, in birds the effects of synchrony in the environment also depended on movement propensity, with a positive impact for resident and short-distance migration species. In insects, annual population synchrony was affected by species movement propensity and dietary niche breadth, but these traits did not modify the effects of environmental synchrony. Our study provides empirical support for the prediction that spatial correlation in population dynamics is more influenced by environmental stochasticity for life histories with lower mobility and faster pace of life, but only in birds. By quantifying spatial population synchrony across different levels of environmental synchrony and life history traits, our study improves the understanding of the Moran effect as well as factors that drive population persistence in the face of environmental change.

## Introduction

Spatial population synchrony, the tendency for geographically separate populations of the same species to exhibit parallel fluctuations in abundance over time, is largely caused by correlated environmental conditions (Liebhold et al., 2004), which typically results in populations closer together in space having more synchronized dynamics (Ranta et al., 1995, Bjørnstad et al., 1999). Studies of spatiotemporal patterns in nature have long relied on the first theory of spatial population synchrony, Moran’s theorem, to explain how the environment causes spatial population synchrony between these spatially separated populations (Moran, 1953, Bjørnstad et al., 1999). Moran’s theorem states that given the same density dependence, populations are expected to show the same synchrony in their population dynamics as the synchrony in their environment (often called the “Moran Effect”; Moran, 1953). When populations are far enough apart for their environments to fluctuate independently of each other, we expect to see no population synchrony (Moran, 1953, Royama, 1992).

Environmental variables significantly affect population dynamics by influencing reproductive success (Lehikoinen et al., 2011, Andreasson et al., 2020), survival rates (Jones et al., 2007, Hansen et al., 2013, Clarke, 2017), immigration rates, and emmigration rates (Pärn & Sæther, 2012). The two most commonly measured environmental variables that have been identified as important drivers of spatial population synchrony are temperature and precipitation (e.g., Post & Forchhammer, 2004, Koenig & Liebhold, 2016, Kahilainen et al., 2018, Dallas et al., 2020, Nicolau et al., 2022), with most results correlating increased synchrony in the environment with increased spatial population synchrony. These variables typically exhibit strong spatial synchrony that declines with distance (Koenig, 2002, Herfindal et al., 2022).

Despite the synchronizing effect of environmental autocorrelation on population dynamics, different species present at the same locations and exposed to the same environmental synchrony do not always exhibit the same degree of synchrony in their population cofluctuations (Marquez et al., 2019, Martin et al., 2023). Different responses to the environment and, thereby, the environmental synchrony are often attributed to life history traits, rendering species-specific sensitivity to changes in the environment (Tedesco & Hugueny, 2006, Chevalier et al., 2014, Hansen et al., 2020). Key life history traits such as position on the fast-slow life history continuum (i.e., an organism’s pace of life derived from generation time or age at first reproduction; Oli, 2004, Gaillard et al., 2005, Reif et al., 2010), movement propensity (i.e., migration classification or distance travelled annually; Howard et al., 2020), and dietary specialization (i.e., the number of food types in the annual diet of a given species; de Gabriel Hernando et al., 2022) are all expected to impact species’ sensitivities to the environment. For example, both theoretical and empirical work shows that environmental stochasticity tends to have a greater effect on population dynamics for species with shorter generation times (Tedesco & Hugueny, 2006, Bjørkvoll et al., 2012, Sæther et al., 2013, Chevalier et al., 2014, Marquez et al., 2019). Distance traveled or migratory tactics are traits that can act as a proxy for a species dispersal ability, which has been shown to strengthen spatial population synchrony (Ranta et al., 1995, Lande et al., 1999, Kendall et al., 2000). Investigating empirically how the Moran effect is modified by such key life history traits is an important next step in understanding the implications of environmental change for spatial population dynamics and, thereby, conservation and the spatial scale of wildlife management actions.

In this study, we compiled a pan-European collection of long-term annual abundance data on birds and insects to identify how species’ life history traits can modify the effects of annual environmental (i.e., temperature and precipitation) synchrony. Birds and insects are informative study organisms for investigating such effects of environmental synchrony on population dynamics because of their history of long-term monitoring and data availability (Nadeau et al., 2017) as well as their large variability in life histories. These taxa are also generally widely distributed, making it possible to study the same species spread across different environments (Jones et al., 2007). Based on Moran’s theorem, we predicted that species of birds and insects in environments with higher synchrony would have overall higher spatial population synchrony, but that the effect of synchronized environments would depend on species’ life history traits (Martin et al. 2023, Marquez et al., 2019). More specifically, we expected that species more sensitive to environmental stochasticity, such as fast-lived species (Sæther et al., 2013), or specialist species (Dumoulin & Armsworth, 2022), would be more highly synchronized and more influenced by environmental synchrony.

## Methods

### i. Bird and insect abundance data

We used population abundance data of breeding birds and insects from eleven long-term monitoring programs located across eight countries: Finland, France, Ireland, the Netherlands, Norway, Sweden, Switzerland, and the United Kingdom (Figure 1A, Table 1). Survey duration was variable, but all were at minimum 10 years long (Table 1). Although there were differences in data collection protocols across countries, as well as between birds and insects, all surveys used either point or line transects, with protocols known for their high quality and rigor (Voříšek et al., 2008, Sevilleja et al., 2020). For bird abundance data, all surveys were conducted during the breeding season, which spanned from spring to mid-summer. For insect abundance data, all surveys were conducted following the Butterfly Monitoring Survey (BMS) standardized protocol of line transects (i.e. fixed routes) repeatedly counted during the butterfly season. These datasets are representative subsets of larger data aggregates (Pan-European Common Bird Monitoring Survey [PECBMS] and Butterfly Monitoring Survey [BMS]; Sevilleja et al., 2020, Brlík et al., 2021). We assumed sampling error was the same across datasets (of birds or insects) because they followed a standardized sampling protocol and were part of a larger consortium of standardized data. Data from these countries were publicly available for download or free to use with data sharing agreements.

**Figure 1.**
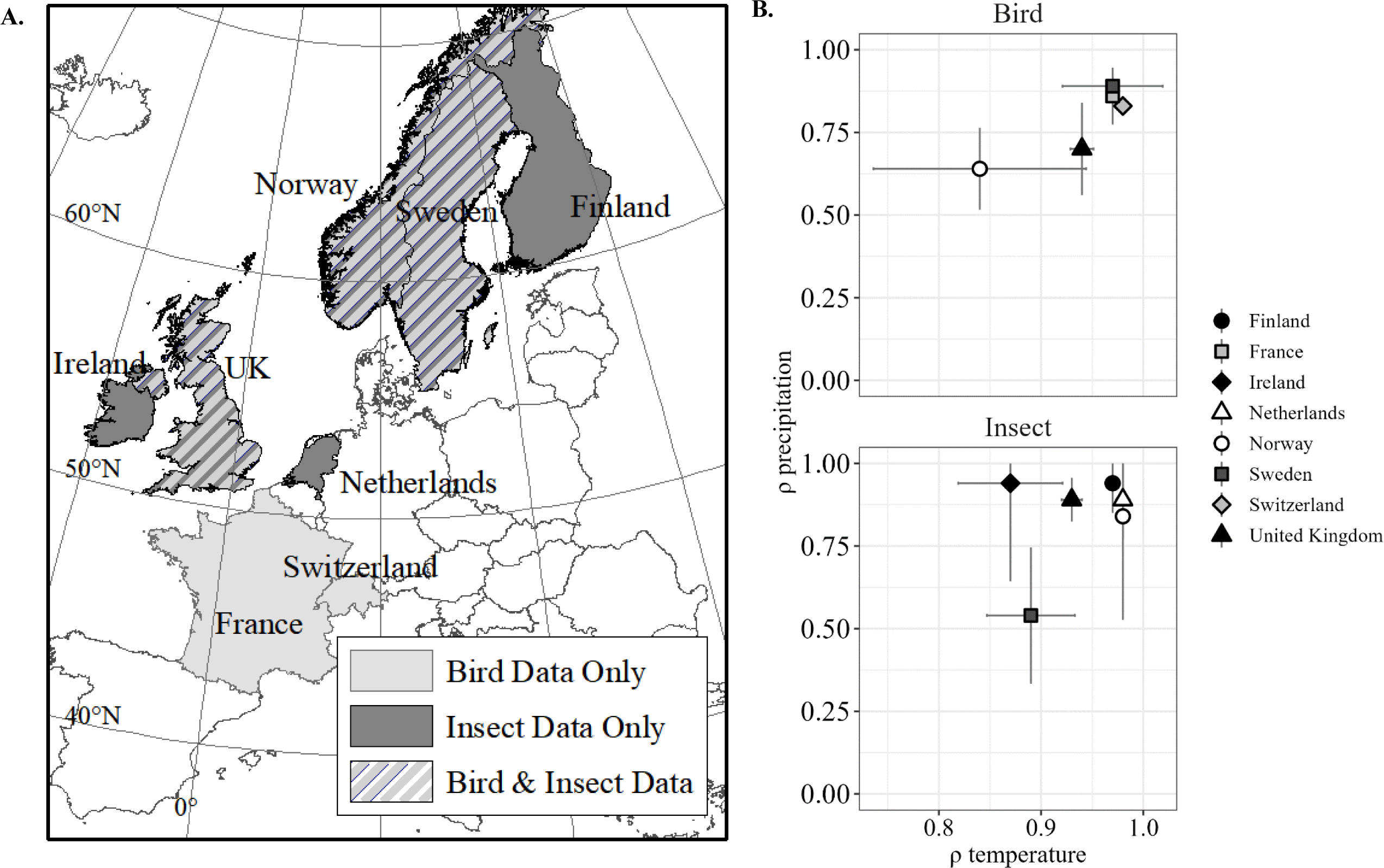
A) Map of European countries from which long-term abundance data were used in this analysis. We analyzed bird data from five countries (France, Norway, Sweden, Switzerland, and the UK [United Kingdom]). We analyzed insect data from six countries (Finland, Ireland, the Netherlands, Norway, Sweden, and the UK). B) Country-specific temperature and precipitation synchrony (*ρ* temperature and *ρ* precipitation) were estimated for bird and insect survey sites.

**Table 1.** Datasets used in analysis. Datasets available either in public domain (e.g., Global Biodiversity Information Service) or through data sharing agreements.

| Country | Survey Name | Taxa | #<br>Species | Years |
| --- | --- | --- | --- | --- |
| Finland | Finnish Butterfly Monitoring Scheme (BMS Finland)<sup>1</sup> | Insects- *Lepidoptera* | 35 | 1999 - 2017 |
| Ireland | Irish Butterfly Monitoring Scheme<sup>2</sup> | Insects- *Lepidoptera* | 15 | 2008 - 2021 |
| Netherlands | The Dutch Butterfly Monitoring Scheme<sup>3</sup> | Insects- *Lepidoptera* | 27 | 1990 - 2020 |
| Norway | Nature Index of Norway: Bumblebees and butterflies in Norway<sup>4</sup> | Insects- *Lepidoptera* & *Bombus* | 10 | 2010 - 2021 |
| Sweden | The Swedish Butterfly Monitoring Scheme (SeBMS)<sup>5</sup> | Insects- *Lepidoptera* & *Bombus* | 40 | 2010 - 2021 |
| United Kingdom | United Kingdom Butterfly Monitoring Scheme (UKBMS)<sup>6</sup> | Insects- *Lepidoptera* & *Bombus* | 37 | 1995 - 2021 |
| France | Suivi Temporel des Oiseaux Communs (STOC)<sup>7</sup> | Birds – Landbirds | 54 | 2001 - 2021 |
| Norway | Norwegian Extensive monitoring of breeding birds (TOV-E)<sup>8</sup> | Birds – Landbirds | 39 | 2006 - 2021 |
| Sweden | Swedish Bird Survey standardrutterna<sup>9</sup> | Birds – Landbirds | 59 | 2006 - 2019 |
| Switzerland | Monitoring Häufige Brutvögel MHB program<sup>10</sup> | Birds – Landbirds | 47 | 1999 - 2020 |
| United Kingdom | BTO/JNCC/RSPB Breeding Bird Survey (BBS)<sup>11</sup> | Birds – Landbirds | 50 | 1994 - 2015 |

<sup>1</sup> Finnish Biodiversity Information Facility (2023). National Finnish butterfly monitoring scheme (NAFI). Occurrence dataset https://doi.org/10.15468/imsrtd accessed via GBIF.org on 30.4.2020.
<sup>2</sup> Liam Lysaght. Irish Butterfly Monitoring Scheme. National Biodiversity Data Centre. Occurrence dataset https://doi.org/10.15468/7wb9as accessed via GBIF.org on 02.06.2023.
<sup>3</sup> The Dutch Butterfly Monitoring Scheme is run by Vlinderstichting (Dutch Butterfly Conservation) and CBS (Statistics Netherlands) as part of the Dutch Network for Ecological Monitoring (NEM).
<sup>4</sup> Åström S, Åström J (2022). Bumblebees and butterflies in Norway. Version 1.4. Norwegian Institute for Nature Research. Sampling event dataset https://doi.org/10.15468/mpsa4g accessed via GBIF.org on 2023-04-12.
<sup>5</sup> Pettersson L B (2022). Swedish Butterfly Monitoring Scheme (SeBMS). Version 1.11. Department of Biology, Lund University. Sampling event dataset https://doi.org/10.15468/othndo accessed via GBIF.org on 2023-04-12.
<sup>6</sup> UK Butterfly Monitoring Scheme (2021). UK Butterfly Monitoring Scheme (UKBMS). Occurrence dataset https://doi.org/10.15468/gmqvmk accessed via GBIF.org on 30.4.2020. Brereton, T.M., Botham, M.S., Middlebrook, I., Randle, Z., Noble D., Harris, S., Dennis, E.B., Robinson A., Peck, K. & Roy, D.B. 2020. UK. Butterfly Monitoring Scheme report for 2019. UK Centre for Ecology & Hydrology, Butterfly Conservation, British Trust for Ornithology and Joint Nature Conservation Committee.
<sup>7</sup> Jiguet F, Devictor V, Julliard R, Couvet D (2012) French citizens monitoring ordinary birds provide tools for conservation and ecological sciences. *Acta Oecologica* 44, 58-66
<sup>8</sup> Kålås J A, Øien I J, Stokke B, Vang R (2021). TOV-E Bird monitoring. Version 1.9. Norwegian Institute for Nature Research. Occurrence dataset https://doi.org/10.15468/qv7d3u accessed via GBIF.org on 2023-04-12.
<sup>9</sup> Lindström Å, Green M (2022). Swedish Bird Survey: Fixed routes (Standardrutterna). Version 1.12. Department of Biology, Lund University. Sampling event dataset https://doi.org/10.15468/hd6w0r accessed via GBIF.org on 2023-04-12.
<sup>10</sup> Swiss monitoring of common breeding birds, Swiss ornithological institute, https://www.vogelwarte.ch/en/projects/monitoring/monitoring-common-breeding-birds.
<sup>11</sup> e BTO Common Birds Census and BTO/JNCC/RSPB Breeding Bird Survey, which provided the data on which these analyses are based. The CBC was funded by the Joint Nature Conservation Committee (JNCC) on behalf of English Nature, Scottish Natural Heritage, Countryside Council for Wales and the Environment and Heritage Service in Northern Ireland, whilst the BBS is jointly funded by the BTO, JNCC and the Royal Society for the Protection for Birds (RSPB).

### ii. Data cleaning and aggregation

We resolved species names across datasets using the Global Names Resolver (*gnr_resolve*) from the *taxize* package for R (Chamberlain S, 2020). Within each country, we aggregated point or transect level count data (hereafter *surveyed sites)* within hexagonal grid cells (hereafter *grid cells)* to represent regional population indices (Appendix 1; Colin et al., 2007). Hexagonal grids are the most appropriate sampling grid for sampling large areas because they reduce bias due to edge effects and have a smaller uniform average distance from the centroid of the grid compared to rectangular grid cells, an important consideration when conducting analyses using distances of grid centroids to one another as done here (Colin et al., 2007). We checked for underlying structure in relation to the size of the grid cell used by running all analyses and comparing results on grid cells with diameters (i.e., distance from one vertex to the opposite vertex) of both 100km and 50km (Appendix 2). Results presented are from grid cells with a diameter of 100km.

For each species separately, we aggregated abundances into a single value representing the sum of abundances in surveyed sites within a given grid cell to mitigate any random fluctuations caused by demographic stochasticity. We analyzed population dynamics at the grid cell level. We divided the total aggregate count of individuals per grid cell by the number of surveyed sites per grid cell to yield an average, which accounted for possible annual variation in the density of sample units (Link & Sauer, 2002). To ensure that only species for which there was sufficient data for synchrony calculations were included in the analysis, we excluded species that were absent from more than 25% of the grid cells that contained survey sites. Also, for each species, we excluded grid cells in which the species was not observed for at least 10 years of the survey duration. After data aggregation and cleaning, we analyzed 126 bird species and 59 insect species.

### iii. Synchrony calculation

We calculated species’ mean spatial population synchrony on log-transformed annual population growth rates (log(*N*_*t*+1_ ⁄ *N*_*t*_)) for each country separately. The strength of the correlation between populations is influenced by directional and temporal trends in their abundance (Loreau & de Mazancourt, 2008). To address these directional trends, spatial population synchrony analyses estimated as the synchrony of population growth rates instead of population abundances (Loreau & de Mazancourt, 2008). This adjustment effectively reduces the influence of changes in population abundance (Tredennick et al., 2017).

The pairwise distance between grid cells at which spatial population synchrony is estimated can change the average calculated synchrony (Pearson & Carroll, 1999, Dungan et al., 2002), with the inclusion of points at large distances reducing the estimation of average synchrony. Therefore, in order to have a standard distance at which we could compare population synchrony across countries, we limited our spatial scale for analysis to pairs of grid cells within 250km of one another. This was the shortest country-specific maximum distance between pairs of grid cells (Switzerland).

In program R (R Core Team 2020), we calculated pairwise Pearson correlations in population growth rates. We Fisher z-transformed these correlations and took the average from pairs of gid cells within 250km of each other. Fisher z-transformation was necessary so that correlations were normally distributed (Silver & Dunlap, 1987). The mean synchrony for each species within each country was then presented as the back transformed mean of the pairwise correlations between all pairs of grid cells within 250km of one another. We measured the distances between grid cells as the Euclidean distances in kilometers from the centroid projected coordinate (EPSG:3035) of grid cell for each pair of cells. Synchrony was only estimated within country, meaning that there were not pairwise correlations across country borders.

### iv. Environmental covariate classification and synchrony estimation

Mean monthly temperatures and mean monthly precipitation were taken from the Climate Research Unit (CRU) at the University of East Anglia (high-resolution gridded 0.5 by 0.5-degree (i.e., approximately 1,700 km^2^ depending on latitude) data of month-by-month variation in climate; Jones, 2022). These data were based on daily or sub-daily observational data from National Meteorological Services and other external agents. We extracted mean monthly environmental covariate values for all grid cells included in the spatial population synchrony analysis. We were only interested in summer season environmental conditions, as the data available were breeding ground abundances. We defined the summer season for each country as the months across the entire study period in which average temperatures for all the grid cells were greater than 5 degrees Celsius, roughly corresponding to the meteorological vegetation growing season (Bootsma, 1994, Linderholm et al., 2008, Körner et al., 2023). Using this approach, each country was allowed different lengths of summer seasons (Appendix 3).

Since population synchrony was analyzed for annual population growth rates, and to reduce effects of shared climate trends on estimates of environmental synchrony among pairs of grid cells, we linearly detrended temperature and precipitation across the years for each dataset and calculated synchrony on these detrended data. For mean annual summer precipitation and mean annual summer temperature separately, we calculated Pearson pairwise correlations between grid cells (Appendix 3). As with spatial population synchrony calculations on the population growth rates, we Fisher z-transformed the correlations and calculated the mean correlation for all grid cells within a 250km distance interval. The mean synchrony for each environmental covariate within each country is presented as the back transformed mean. We checked for correlations between temperature and precipitation at bird and insect surveyed sites using cross correlations.

### v. Life history trait classification

We characterized each bird or insect species using a range of species-specific traits: position on the fast-slow life history continuum (generation time for birds, voltinism for insects), movement propensity (migratory tactic for birds, months in flight for insects), and specialist/generalist species (dietary diversity for birds and larval dietary breadth for insects; Table 2, Figure 2). We checked for dependencies or correlations between life history traits used in the analysis using Chi-square test of independence for categorical variables, ANOVA for categorical and continuous variables, and cross correlations for continuous variables (Appendix 4).

**Figure 2.**
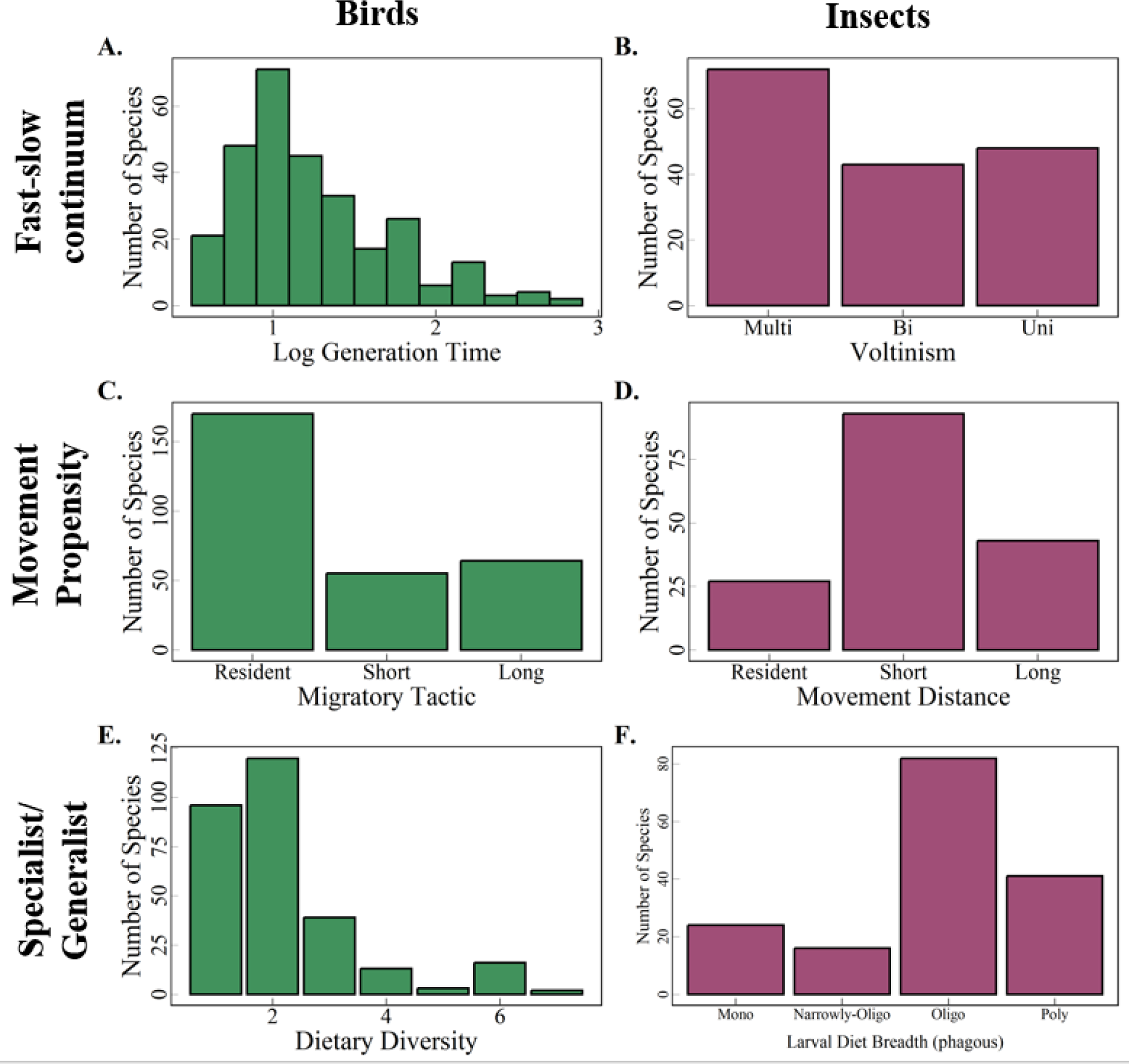
Distributions of the life history traits for birds (in green) and insects (in purple) are presented. (A,B) The number of species classified along the fast-slow life history continuum is presented: we used log generation time for birds, and we used voltinism for insects. Voltinism is reordered so that increasing along the voltinism axis is equivalent to increasing from fast-(multi) to slow-lived (uni) species. Log generation time ranged from 0.53 (absolute scale: 1.69 yr.) to 3.1 (absolute scale: 22.1 yr.). (C,D) The number of species classified according to their movement propensity: We used migratory tactic for birds, and we used movement distance for insects. (E,F) The number of species classified according to the specialization of their diet: We used dietary diversity for birds and we used larval diet breadth for insects. For definitions of life history traits used, see Table 2.

**Table 2.** Life history trait variables used in analysis. Definitions and any modifications to the variables taken from its original source are explained.

| **Taxa** | **Life History Trait** | **Category** | **Definition** | **Modifications** | **Categorical/**<br>**Continuous** | **Range/Categories** |
| --- | --- | --- | --- | --- | --- | --- |
| Birds | Log generation time<sup>12</sup> | Fast-slow | The average age of parents. Bird et al. (2020) classified birds worldwide using age of first reproduction, maximum longevity, and annual adult survival to derive generation times. | Log transformed generation time | Continuous | 1.7 - 25.3 yr. (0.53 - 3.23 on log scale) |
| Insects | Voltinism<sup>13</sup> | Fast-slow | The number of generations each year. | N/A | Categorical | U = Univoltine, B = Bivoltine, M= Multivoltine |
| Birds | Migratory Tactic<sup>13</sup> | Movement Propensity | Resident species were defined as non-migrants with no seasonal movements beyond their country of residence Short-distance migrants had documented non-breeding areas within Europe but outside the country of their breeding ground. Long-distance migrants had documented non-breeding areas outside of Europe. | N/A | Categorical | Resident, Short-distance migrant, Long-distance migrant |
| Insects | Movement Distance<sup>14</sup> | Movement Propensity | The number of months in a given year in which mobile. | Transformed from continuous range of 1-12 months into 3 distinct categories: Resident species (moved for 1- 4 months of a year), short movement (moved for 5 - 8 months of a year), and longer movement distance species (moved for 9 - 12 months of a year). | Categorical | Resident, Short-distance migrant, Long-distance migrant |

<sup>12</sup> Bird, J. P., R. Martin, H. R. Akçakaya, J. Gilroy, I. J. Burfield, S. T. Garnett, A. Symes, J. Taylor, Ç. H. Şekercioğlu, and S. H. M. Butchart. 2020. Generation lengths of the world's birds and their implications for extinction risk. *Conservation Biology* 34:1252-1261.
<sup>13</sup> *Lepidoptera* species: Shirey, V., E. Larsen, A. Doherty, C. A. Kim, F. T. Al-Sulaiman, J. D. Hinolan, M. G. A. Itliong, M. A. K. Naive, M. Ku, M. Belitz, G. Jeschke, V. Barve, G. Lamas, A. Y. Kawahara, R. Guralnick, N. E. Pierce, D. J. Lohman, and L. Ries. 2022. LepTraits 1.0 A globally comprehensive dataset of butterfly traits. *Scientific Data* 9:382. For *Bombus* species: Løken, A. 1973. Studies on Scandinavian Bumble Bees (Hymenoptera, Apidae). *Norsk ent. Tidsskr.* 20, 1–218., pers. comm Sondre Dahle, March 28, 2023.

|  |  |  |  |  |  |  |
| --- | --- | --- | --- | --- | --- | --- |
| Birds | Dietary Diversity<sup>14</sup> | Specialist/<br>Generalist | Nr. of food types composing $\geq 10\%$ of the annual diet. Food type categories were no data, leaves, fruit, grains, arthropods, other invertebrates, fish, other vertebrates, or carrion. | N/A | Continuous | 1 - 9 |
| Insects | Larval Dietary Breadth<sup>14</sup> | Specialist/<br>Generalist | Diversity of larval diet breadth. Monophagous species (1) were at the specialist end of the spectrum, while multiphagous species (4) were at the generalist end of the spectrum. | Transformed categorically increasing scale from<br>M=Monophagous,<br>NO= Narrowly Oligophagous, O= Oligophagous, MP = Multiphagous to continuous scale 1-4 | Continuous | Increasing scale from 1=Monophagous to 4=Multiphagous |

<sup>14</sup> Storchová, L., and D. Hořák. 2018. Life-history characteristics of European birds. *Global Ecology and Biogeography* 27:400-406.

*Fast-slow life history:* We used generation time as a proxy for classification of bird species along the fast-slow life history continuum (Gaillard et al., 2005, Bjørkvoll et al., 2012, Martin et al., 2023). The fast-slow life history continuum ranges from species with short generation times that are fast-reproducing and short-lived (i.e., fast-lived) to species with long generation times that are slow-reproducing and long-lived (i.e., slow-lived; Stearns, 1983; Gaillard et al., 1989; Galliard et al., 2016). In this study, we used species-specific generation times from Bird et al. (2020), who derived generation times for birds worldwide using proxies such as the age of first reproduction, maximum longevity, and annual adult survival. We log transformed generation time for use in the analysis.

We classified each insect species along the fast-slow life history continuum using voltinism (i.e., the number of generations of species each year; for *Lepidoptera* species: Shirey et al. 2022, for *Bombus* species: Løken, 1973, pers. comm Sondre Dahle). Voltinism has been used to explain insects’ degree of vulnerability to climatic events (e.g., Melero et al., 2016) and is a useful proxy for position on the fast-slow life history continuum in species that do not have readily available generation time information (Kőrösi et al., 2022). Fewer generations per year (i.e., univoltine) are associated with a slower-lived species, whereas more generations per year (i.e., multivoltine) are associated with faster-lived species.

#### Movement propensity

We classified each bird species as a resident, short-distance, or long-distance migrant (following Martin et al. 2023). Avian species that migrate are usually categorized based on the extent of their movement between breeding and overwintering regions (Rappole, 2013). In this study, resident species were defined as those that remained in their country of residence throughout the year, without undertaking seasonal movements (Newton, 2008, Eyres et al., 2017). Species considered short-distance migrants were those that has documented non-breeding areas within Europe but outside the country of their breeding ground (Rappole, 2013). Long-distance migrants were those species that had documented non-breeding areas located outside of Europe (Rappole, 2013). We used an available database of avian life history traits (Storchová & Hořák, 2018) to classify each bird species into one of the three migration tactics, i.e., residents, short-distance migrants, or long-distance migrants.

We classified each insect species according to their movement distance: Insects could have long-distance movement, short-distance movement, or have ‘no’ movement based on their flight duration (i.e., the number of months each year in which species were mobile; for *Lepidoptera* species: Shirey et al. 2022, for *Bombus* species: Løken 1973, pers. comm Sondre Dahle). Here, movement distance and flight duration can be considered a proxy for insect migration distance and can provide valuable information about an insect’s capability and propensity for long-distance travel. Insect migration differs from bird migration in that insects rarely complete annual circular movements between breeding and non-breeding grounds, and most movements require multiple generations to complete (Chapman et al. 2015). Following Dingle and Drake (2007), we therefore defined insect migration as the persistent, straightened-out movement typically carrying an individual away from a location where they were produced to another where they breed (Dingle & Drake, 2007). This persistent movement can be quantified as the amount of time in which a species is in flight (i.e., flight duration; Minter et al., 2018), or the distance traveled (i.e., flight distance). The two are correlated (Guo et al., 2020). We transformed the flight duration data from a continuous range of 1-12 months into 3 distinct categories: Resident species (species that moved for 1-4 months of a year), short movement species (species that moved for 5 - 8 months of a year), and longer movement species (species that moved for 9 - 12 months of a year). One may expect that these two types of movement (bird migration and insect movement) would have a similar impact on spatial population synchrony, since, for both taxa, we assume time spent moving was time spent away from a shared breeding ground, which may have acted to disrupt synchrony.

#### Specialist/generalist

We classified each bird species along a continuum of one (specialist) to nine (generalists) according to their dietary diversity (i.e., breadth). The value used in the analysis corresponded to the total number of different food types in the annual diet of a given species (Storchová & Hořák, 2018). Species could be classified as eating leaves, fruit, grains, arthropods, other invertebrates, fish, other vertebrates, or carrion. A species was recorded as eating a type of food if that food comprised at least 10% of its diet throughout the year (Storchová & Hořák, 2018).

We classified each insect species according to their larval diet breadth using a global lepidoptera trait database (Kőrösi et al., 2022). Larval diet breadth was a categorical variable with monophagous species, or species that ate only one kind of food, at the specialist end of the spectrum, with polyphagous species, or species that ate multiple kinds of food, at the generalist end of the spectrum (for *Lepidoptera* species: Kőrösi et al., 2022, for *Bombus* species: Løken 1973, pers. comm Sondre Dahle).

### vi. Evaluating the impact of environmental synchrony and life history traits on synchrony in annual population growth rates

We used linear mixed models on bird and insect data separately to determine if there was an effect of environmental synchrony on spatial population synchrony across species, while accounting for life history traits. Given the collinearity between synchrony in temperature and precipitation at both bird and insect surveyed sites (correlation of 0.86 and 0.89 respectively), we built two different model sets to test the effect of these environmental covariates independent from one another. We included models that included an interaction between the environmental covariate and life history traits to determine if species had different responses to environmental synchrony depending on trait differences. In our separate global models for birds and insects (Table 3), we included species as a random effect and added position on the fast-slow life history continuum (continuous), movement propensity (categorical), specialist/generalist (continuous), mean synchrony in temperature or mean synchrony in precipitation as fixed effects. To account for potential bias in the distribution of survey points, we also included a covariate in the models that represented the median distance between populations at which spatial population synchrony was estimated. We used Akaike information criterion adjusted for small sample size (AICc) based on models fitted with maximum likelihood (ML) to rank models (Burnham & Anderson, 2002, Bolker et al., 2009). Parameter estimates and their uncertainties were based on models fitted with restricted maximum likelihood estimators (REML). Residuals were checked for normality.

**Table 3.** Model selection results for the analysis of spatial synchrony in annual population growth rates of (A) birds and (B) insects in Europe. Synchrony estimates are based on pairs of populations *≤*250km apart, merged in grid cells of size 100km diameter. Covariates designated with a “+” were present in model. Covariates included environmental synchrony (Env; in terms of mean summer precipitation [Precip] or mean summer temperature [Temp]), movement (Mvmt), fast-slow life history continuum (Fastslow), specialist/generalist (Specgen), and two-ways interactions between environmental synchrony and the life history traits. We also included a covariate for median distance at which synchrony was calculated (Med dist). Only one environmental covariate (precipitation or temperature) was included in each model because of collinearity, resulting in two different model sets. Only the top five models are presented (rank 1-5). We relied upon Akaike’s Information Criterion with a small sample size correction (*AICc)* for model selection and used Akaike model weights (*wt*) and *ΔAICc* to identify the top model. Number of parameters in model indicated by column k. LogLik = log-likelihood.

| (A) Birds |  |  |  |  |  |  |  |  |  |  |  |  |  |  |  |
| --- | --- | --- | --- | --- | --- | --- | --- | --- | --- | --- | --- | --- | --- | --- | --- |
| Taxa | Rank | Env. Variable | Env | Mvmt | Fastslow | Specgen | Mvmt x Env | Fastslow x Env | Specgen x Env | Med dist | k | LogLik | AICc | $\Delta AIC_c$ | $wt$ |
| **Birds** | **1** | Precip | + | + | + | -- | + | -- | -- | + | **11** | **217.92** | **-412.89** | **0** | **0.13** |
| Birds | 2 | Precip | -- | + | + | -- | -- | -- | -- | + | 8 | 214.54 | -412.58 | 0.31 | 0.11 |
| Birds | 3 | Precip | + | + | + | -- | + | + | -- | + | 12 | 218.52 | -411.92 | 0.97 | 0.08 |
| Birds | 4 | Precip | + | + | + | + | + | -- | -- | + | 12 | 218.16 | -411.19 | 1.70 | 0.06 |
| Birds | 5 | Precip | + | + | + | -- | + | -- | -- | -- | 10 | 215.92 | -411.05 | 1.84 | 0.05 |
| **Birds** | **1** | Temp | + | + | + | -- | + | + | -- | -- | **11** | **219.84** | **-416.72** | **0** | **0.16** |
| Birds | 2 | Temp | + | + | + | + | + | + | + | + | 12 | 221.53 | -415.75 | 0.33 | 0.14 |
| Birds | 3 | Temp | + | + | + | + | + | + | -- | -- | 13 | 220.19 | -415.25 | 0.97 | 0.09 |
| Birds | 4 | Temp | + | + | + | -- | -- | + | -- | -- | 14 | 216.16 | -413.67 | 1.00 | 0.07 |
| Birds | 5 | Temp | + | + | + | + | -- | + | + | + | 12 | 218.12 | -413.29 | 1.47 | 0.06 |

**(B) Insects**
| Taxa | Rank | Env. Variable | Env | Mvmt | Fastslow | Specgen | Mvmt x Env | Fastslow x Env | Specgen x Env | Med Dist | k | LogLik | AICc | $\Delta AIC_c$ | wt |
| --- | --- | --- | --- | --- | --- | --- | --- | --- | --- | --- | --- | --- | --- | --- | --- |
| **Insects** | **1** | Precip | + | + | -- | + | -- | -- | -- | -- | **7** | **88.82** | **-162.93** | **0** | **0.18** |
| Insects | 2 | Precip | + | + | + | + | -- | -- | -- | -- | 9 | 90.83 | -162.51 | 0.42 | 0.14 |
| Insects | 3 | Precip | + | -- | + | + | -- | -- | -- | -- | 7 | 88.51 | -162.31 | 0.63 | 0.13 |
| Insects | 4 | Precip | + | + | -- | + | -- | -- | + | -- | 8 | 89.21 | -161.49 | 1.44 | 0.09 |
| Insects | 5 | Precip | + | + | + | + | -- | -- | + | -- | 10 | 91.26 | -161.09 | 1.83 | 0.07 |
| **Insects** | **1** | Temp | + | + | -- | + | -- | -- | -- | -- | **7** | **61.19** | **-107.67** | **0** | **0.15** |
| Insects | 2 | Temp | + | + | -- | + | -- | -- | -- | -- | 9 | 63.12 | -107.08 | 0.59 | 0.11 |
| Insects | 3 | Temp | -- | + | -- | + | -- | -- | -- | -- | 6 | 59.55 | -106.56 | 1.11 | 0.09 |
| Insects | 4 | Temp | + | + | + | + | -- | -- | -- | -- | 9 | 62.69 | -106.21 | 1.46 | 0.07 |
| Insects | 5 | Temp | + | + | -- | + | -- | -- | + | -- | 8 | 61.41 | -105.89 | 1.77 | 0.06 |

## Results

Of the 126 unique bird species analyzed, 13 species were present in five countries, 14 in four countries, 16 in three countries, 36 in two countries, and 47 in one country. Of the 59 unique insect species analyzed, six species were present in six countries, ten in five countries, ten in four countries, ten in three countries, seven in two countries, and 16 in one country. Average synchrony across all insect species was 0.31 (SD=0.03), while average synchrony across all bird species was 0.09 (SD=0.01; Figure 3B). Estimates of spatial population synchrony were thus generally higher for insects than for birds (Figure 3A). For bird and insect data present in the same country (i.e., in Norway, Sweden, and United Kingdom), insects had higher mean synchrony (Figure 3A). For species-specific estimates of synchrony, see Appendix 5.

**Figure 3.**
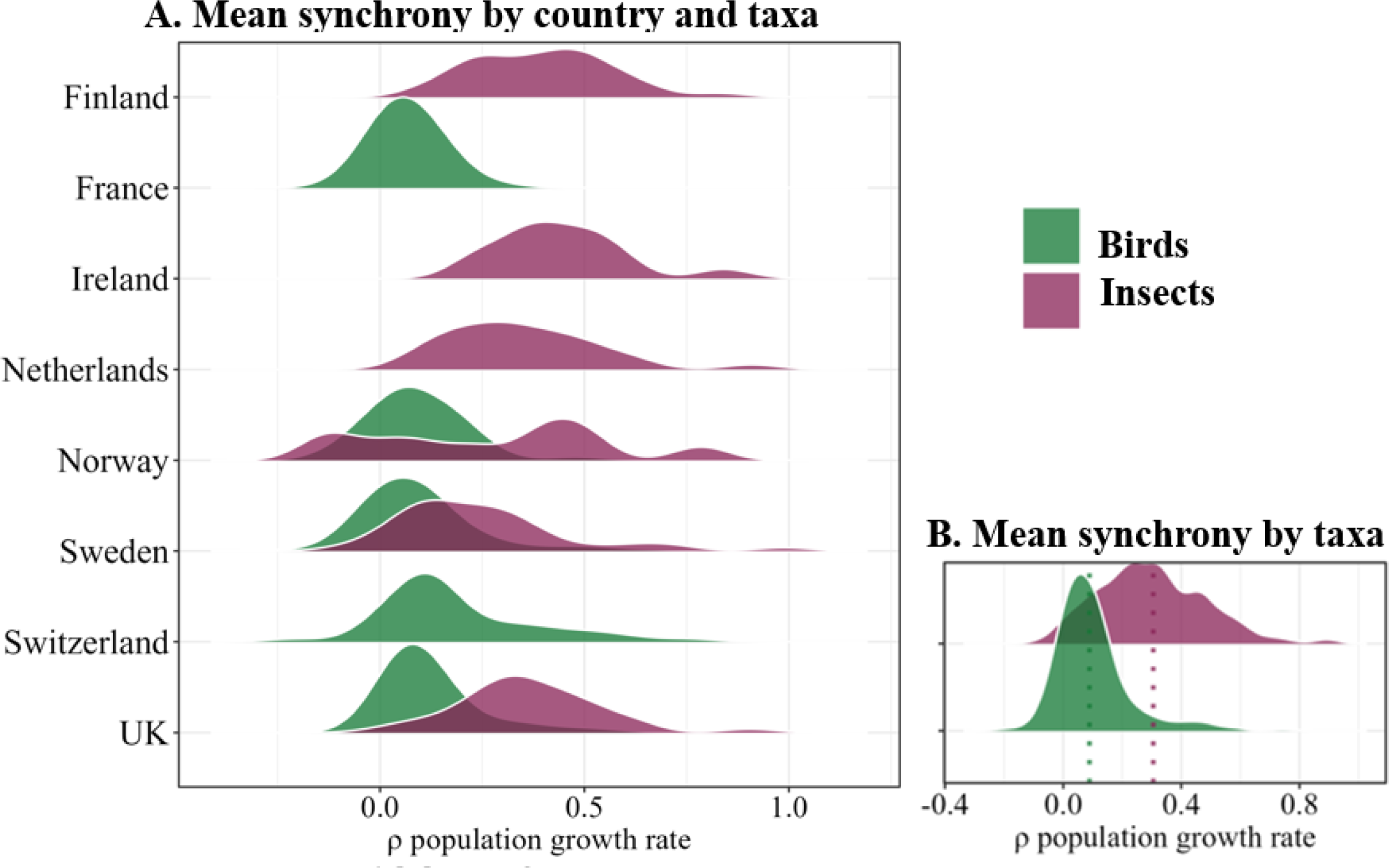
A) Distributions of the estimates of mean spatial synchrony in annual population growth rates for species of birds and insects, separated by country (birds shown in green, insects shown in purple). Distributions of mean synchrony are calculated from the R package *ggridges* function *geom_density_ridges*, which computes a kernel density estimate from the data. B) Distributions of the estimates of mean synchrony when combined across all countries are shown. Mean synchrony estimates are indicated by dotted line per taxonomic group.

There was strong support for several of the top candidate models in our model sets for insects and birds (ΔAICc < 2.0; Table 3). The synchronizing effect of the environment, either as precipitation or temperature, was present in 9 out of ten top models for birds (ΔAICc < 1.84; Table 3A) and in 9 out of ten top models for insects (ΔAICc < 1.83; Table 3B). There was strong evidence that there was an environmental effect driving spatial population synchrony across the datasets analyzed.

For birds, there was strong support for an effect of environmental synchrony on population synchrony (Table 3; Figure 4A-C), and this synchronizing effect of the environment depended on life history traits. For birds, the highest ranked model which did not include an interaction between a life history trait and temperature was ranked twelfth and had a ΔAICc = 4.13, whereas the highest ranked model which did not include an interaction between a life history trait and precipitation was ranked second and had a ΔAICc = 0.31 (Table 3). Temperature had a stronger synchronizing effect than precipitation (Figure 4A-C, Table 3). The model with the strongest support indicated that synchrony in population growth rates increased with increasing synchrony in temperature, but only for short distance migrants (β=1.61, SE=0.34) and resident species (β=1.63, SE=0.51) compared to long-distance migrants (β=0.75, SE=0.36). Moreover, as synchrony in temperature increased, species with shorter generation times showed a larger increase in synchrony than species with longer generation times (Figure 4A). Regarding precipitation, the highest ranked model indicated that synchrony in population growth rate in birds was explained by synchrony in precipitation, movement propensity, and position on the fast-slow life history continuum (Table 3, Table 4). The effect of synchrony in precipitation depended on a species’ movement propensity (resident species: β=0.29, SE=0.10, short-distance migrant: β=0.31, SE=0.12, long-distance migrant: β=0.18, SE=0.18; Figure 4C). Resident species and short-distance migrants were positively impacted by increasing synchrony in temperature and precipitation (Figure 4B-C). Position on the fast-slow life history continuum was an important predictor of spatial population synchrony but did not interact with synchrony in precipitation (Figure 4D).

**Figure 4:**
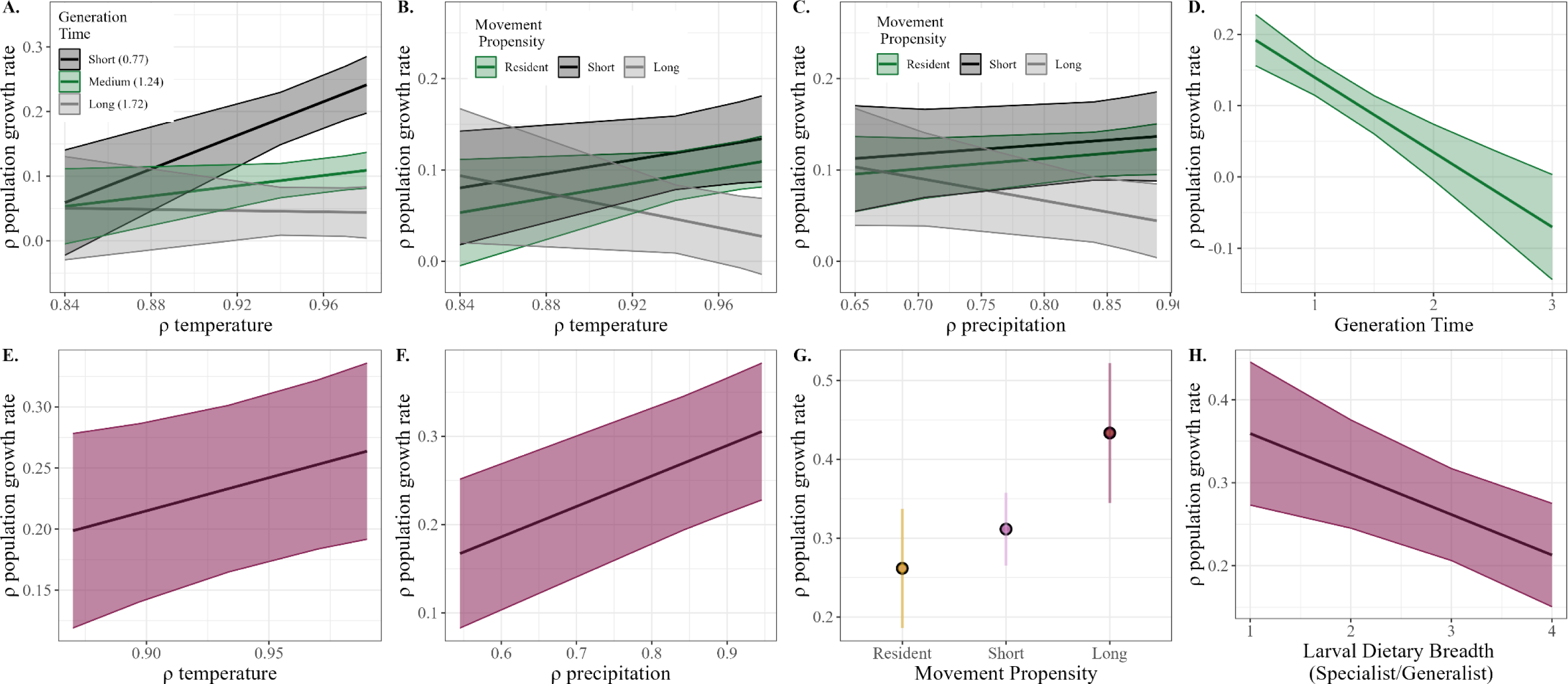
Effects of environmental variables and life history trait covariates included in top models for (A-D) birds and (E-F) insects. A) Synchrony in temperature and generation time in birds. B). Synchrony in precipitation and movement propensity in birds. C). Synchrony in temperature and movement propensity in birds. D) The effect of generation time on population growth rate in birds. E) Synchrony in temperature and spatial population growth rate in insects. F) Synchrony in precipitation and synchrony in population growth rates in insects. G) The effect of movement propensity on synchrony in population growth rate in insects. H) The effects of larval dietary breadth on synchrony in population growth rate in insects. 95% confidence intervals presented as shaded colors.

**Table 4.** Parameter estimates, standard errors, and 95% confidence intervals (CI) for the (A) bird and (B) insect top models from the model selection of spatial synchrony in annual population growth rates. Interactions indicated between variables with an “x”.

**Temperature**
| Parameter | Estimate | Std. Error | 95% CI |
| --- | --- | --- | --- |
| Mean synchrony in temperature | 1.63 | 0.51 | 1.61 – 2.65 |
| Resident species | -1.32 | 0.48 | -2.28 – -0.36 |
| Short-distance migration | -1.28 | 0.40 | -2.08 – -0.48 |
| Long-distance migration | -0.55 | 0.45 | -1.45 – 0.35 |
| Fast-slow life history continuum | 0.76 | 0.31 | 0.14 – 1.38 |
| Mean synchrony in temperature x<br>Fast-slow life history continuum | -0.91 | 0.32 | -1.55 – -0.27 |
| Mean synchrony in temperature x<br>Short-distance migrants | -0.02 | 0.34 | -0.69 – 0.66 |
| Mean synchrony in temperature x<br>Long-distance migrants | -0.88 | 0.36 | -1.59 – -0.15 |
| $r^2$ marginal | $r^2$ conditional | | |
| 0.17 | 0.45 |  |  |

**Precipitation**
| Parameter | Estimate | Std. Error | 95% CI |
| --- | --- | --- | --- |
| Mean synchrony in precipitation | 0.11 | 0.10 | -0.09 – 0.31 |
| Resident species | 0.18 | 0.09 | -0.01 – 0.37 |
| Short-distance migration | 0.21 | 0.12 | -0.03 – 0.45 |
| Long-distance migration | 0.43 | 0.13 | -0.03 – 0.69 |
| Fast-slow life history continuum | -0.11 | 0.02 | -0.15 – -0.07 |
| Mean synchrony in precipitation x<br>Short-distance migrants | -0.01 | 0.17 | -0.36 – 0.32 |
| Mean synchrony in precipitation x<br>Long-distance migrants | -0.36 | 0.18 | -0.72 – 0.00 |
| Median distance | -0.00 | -0.00 | *na* |
| $r^2$ marginal | $r^2$ conditional | | |
| 0.14 | 0.38 |  |  |

**(B) Insects:****Temperature**
| **Parameter** | **Estimate** | **Std. Error** | **95% CI** |
| --- | --- | --- | --- |
| Mean synchrony in temperature | 0.54 | 0.27 | 0 – 1.08 |
| Movement propensity: Resident | -0.13 | 0.27 | -0.67 – 0.41 |
| Movement propensity: Short | 0.10 | 0.26 | -0.42 – 0.62 |
| Movement propensity: Long | 0.03 | 0.26 | 0.04 – 0.26 |
| Specialist/Generalist | -0.05 | 0.02 | -0.08 – -0.01 |
| $r^2$ marginal | $r^2$ conditional | | |
| 0.26 | 0.45 |  |  |

**Precipitation**
| **Parameter** | **Estimate** | **Std. Error** | **95% CI** |
| --- | --- | --- | --- |
| Mean synchrony in precipitation | 0.35 | 0.06 | 0.23 – 0.47 |
| Movement propensity: Resident | 0.13 | 0.09 | -0.05 – 0.31 |
| Movement propensity: Short | 0.18 | 0.09 | 0 – 0.36 |
| Movement propensity: Long | 0.30 | 0.08 | 0.14 – 0.46 |
| Specialist/Generalist | -0.05 | 0.02 | -0.09 – -0.01 |
| $r^2$ marginal | $r^2$ conditional | | |
| 0.35 | 0.56 |  |  |

There was also strong support for an effect of environmental synchrony in insects (Table 3; Figure 4E-F). For insects, the synchronizing effect of precipitation had a stronger effect than temperature (precipitation: β=0.35, SE=0.06, temperature: β=0.54, SE=0.27; Figure 4E-F), but both covariates were in the highest ranked models of their respective model sets (Table 3). For insects, there was weak support for that life history traits influenced the strength of the effect of environmental synchrony on population synchrony. The highest ranked model which included an interaction between a life history trait and temperature was ranked fifth and had a ΔAICc = 1.77, whereas the highest ranked model which included an interaction between a life history trait and precipitation was ranked fourth and had a ΔAICc = 1.44. The model which had the most support across model sets indicated that synchrony in population growth rate was explained best by synchrony in precipitation (β=0.35, SE=0.06), movement propensity (resident species: β=-0.13, SE=0.09, short-distance movement: β=0.18, SE=0.09, long-distance movement: β=0.30, SE=0.08), and classification as specialist/generalist (β=-0.05, SE=0.02; Table 3, Figure 4G-H), but no interaction between life history trait and precipitation synchrony (Table 3). The highest ranked temperature model gave the same top model, but there was weak support for all variables included (Table 3B). Synchrony in population growth rate was explained by synchrony in temperature (β=0.54, SE=0.27), movement propensity (resident species: β=-0.13, SE=0.27, short-distance movement: β=0.10, SE=0.26, long-distance movement: β=0.03, SE=0.26), and specialist/generalist, with increasing degree of generalization resulting in decreased synchrony (β=-0.05, SE=0.02; Table 3).

Despite not being in the top model, there was also support for the inclusion of specialist/generalist classification in interaction with temperature among the bird model sets (ΔAICc = 0.33; Table 3A). For insects, there was support for an effect of fast-slow life history traits on spatial population synchrony (ΔAICc = 0.42; Table 3B) that was evident in all models except the top model.

We confirmed that there was some underlying structure in the data by conducting the analysis on grid cells with a diameter of 50km in addition to the grid cell diameter of 100km presented here (Appendix 2). The top models with 50km diameter grid cells resulted in a few parameter changes in the top models. For birds, the top models no longer included interactions between environmental synchrony and life history traits. However, the main strong effects of generation time and an environmental variable were still present (Appendix 2). For insects, there were fewer differences. The strong main effects of specialist/generalist classification and environmental synchrony were present in all top models for insects (Appendix 2). The loss of the interaction term with analysis at the 50km grid cell size is likely because the total number of surveyed sites within a 50km grid cell were few in some countries (e.g., Sweden averaged 1.2 survey point per 50km grid cell), adding noise to the estimates of population abundance. We also confirmed that there was no spatial bias with respect to how pairs of grid cells were distributed in space on the estimates of synchrony by testing for an effect of median distance at which synchrony was calculated (Appendix 6).

## Discussion

Here, based on datasets of annual abundances of European birds and insects, we advance the empirical understanding of spatiotemporal population dynamics by showing that variation in the impacts of environmental synchrony on spatial population synchrony can depend on the species’ life history traits. In both birds and insects, we found strong evidence that spatial synchrony in precipitation and/or temperature had a positive effect on annual spatial synchrony in population growth rates, indicating a Moran effect (Figure 4). Although synchrony in temperature and precipitation was highly correlated, population synchrony in birds appeared more strongly influenced by temperature than precipitation, and vice versa in insects (Table 3). In birds, the strength of the Moran effect depended on key life history traits. More specifically, responses to increased environmental synchrony depended on generation time and movement propensity, with a positive impact found only for short generation times (i.e., ‘fast’ species) and for resident and short-distance migration species (Figure 4). In contrast, for insects, movement propensity and dietary niche breadth influenced population synchrony but, at the temporal scale investigated here, these or other life history traits did not appear to modify the overall positive effect of environmental synchrony.

Although we do not demonstrate causality here, the synchronized environmental factors likely had a synchronizing effect on population dynamics via the Moran effect (Moran, 1953). Synchrony in the environment, either temperature or precipitation, was high at survey sites ≤ 250km apart (Figure 1B, Appendix 3). Synchrony in temperature was higher than synchrony in precipitation across all countries except Ireland, matching previously identified relationships between precipitation and temperature (e.g., Koenig, 2002, Herfindal et al., 2020). For many species, the environment experienced during the spring and breeding season is particularly important for driving fluctuations in parameters of importance to lifetime fitness and survival (Crick, 2004, Pearce-Higgins et al., 2015). Environmental conditions such as average summer precipitation and average summer temperature are known to act as important constraints on population growth rates of both birds and insects (Crick, 2004, Zipkin et al., 2012, Pearce-Higgins et al., 2015, Meller et al., 2018, Herrando et al., 2019). In this paper we have documented the same effects of the environment in two quite different taxonomic groups, indicating general patterns relevant at large spatial scales.

We found support for life history traits in interaction with the environment in birds, meaning that different groups of species responded differently to environmental synchrony. As far as we are aware, this is the first time interactions between life history traits and environmental synchrony have been documented to impact spatial population synchrony. Our results add knowledge about spatial population synchrony by showing that species with certain traits are more likely to respond to synchrony in the environment. Empirically, we have shown the importance of considering a species’ life history traits when predicting the impacts of the environment on spatial population synchrony.

We further extend what is known about the importance of temperature to avian population dynamics by including the interaction effect with life history traits. Temperature during the breeding season interacted with avian position on the fast-slow life history continuum. Generally, for birds, species with shorter generation times had higher synchrony in population growth rates. There was no notable effect of increasing synchrony in the environment for species with long generation times, suggesting that they are less sensitive to environmental conditions. These general patterns match what is expected based on theory. Theoretical and empirical examples show that environmental stochasticity has a greater effect on population dynamics for species with shorter generation times, which tend to have more immediate responses to environmental stochasticity (Sæther et al., 2005). For example, Sæther et al. (2013) found that the stochastic influence of the environment on population dynamics of a species decreased as generation time increased, resulting in decreased overall stochasticity of population dynamics.

For movement propensity in birds, higher synchrony in temperature and precipitation was associated with higher spatial population synchrony for resident and short-distance migrants. Long-distance migrants had lower synchrony with increasing environmental synchrony. We expected to find the highest effect of synchrony in the breeding ground environment for resident species because two resident populations are more likely to experience the same or similar seasonal changes in environmental conditions for a longer duration than migrant or nomadic species, which typically spend less time on the breeding grounds. Migrants can spend as few as four months on the breeding ground before departing for wintering grounds (e.g., long-distance migrants; Knaus et al., 2018). Patterns of synchrony for short-distance migrants mirror the patterns identified in resident species. This could be occurring because short-distance migrants by our classification schema migrated within Europe, meaning that the over-wintering grounds they went to could still have environments which were synchronized with the breeding ground environmental dynamics (Butler, 2003).

Independent from environmental synchrony, insect species’ life history traits were strong predictors of spatial population synchrony. Despite finding evidence for an interaction between environment and life history traits in birds, we found little support for the same interaction in insects. It is possible such an interaction exists on smaller temporal or spatial scale, and that the scales used to measure environmental synchrony in this study was too large for the scale of insect life cycles (Jan et al., 2017). Further studies testing for an interaction between life history traits and environmental synchrony in insects should consider looking at varying temporal and spatial scales. Studies which have investigated average daily temperature and precipitation during insect flight season (typically summer) found that both impacted spatiotemporal dynamics of butterfly species (Gibbs et al., 2011).

For insects, movement propensity was an important predictor of spatial population synchrony. Species that were resident or characterized by short-distance movement had similar spatial population synchrony, which was lower than spatial population synchrony of species characterized by long-distance movement. While this is not the result we would expect if the short-distance and long-distance movement species were true “migrants”, this is the expected result if long-distance movement can also encompass movement by dispersal. Insect movement is classified here as number of months a species is in flight and is expected to follow the classical theory of dispersal driving spatial population synchrony (Lande et al., 1999, Ims & Andreassen, 2005). With increased dispersal, increased synchrony occurs as individuals from a population at high density move to a population with lower density, resulting in a smaller difference in density between the two populations (Ripa, 2000). Finally, specialist species were more synchronized than generalist species. Specialist species have known higher sensitivity to environmental stochasticity than generalist species (Dumoulin & Armsworth, 2022), but linking this to spatial population synchrony has rarely been shown empirically.

While both temperature and precipitation were important predictors for the annual population synchrony in birds and insects, we found strong support showing that temperature is the more important of the two environmental variables for synchronizing bird dynamics and that precipitation is more important for synchronizing insect dynamics. Others have found that summer precipitation synchronized population dynamics (regardless of life history strategy) in *Lepidoptera* species (e.g., Glanville fritillary butterfly (*Melitaea cinxia*), a species included in this analysis; Kahilainen et al., 2018). Late spring and/or early summer precipitation is known to be important for insects as a trigger for the end of diapause (i.e., a state of arrested development), and for subsequent larval host-plant production (Wolda, 1988). The different important positively synchronizing variables for birds and insects extend the finding of Pearce-Higgins et al. (2015), who found a positive relationship between the mean effect of temperature and population size for birds, but not invertebrates, suggesting that temperature played a larger role in population dynamics for birds. Other studies have linked declining synchrony in temperature to declining bird population synchrony (Koenig, 2001, Koenig & Liebhold, 2016).

The higher spatial population synchrony we identified across bird and insect species in more synchronized environments has implications for future population stability and species persistence under climate change and intensified human use scenarios (Møller et al., 2004). Understanding general patterns in the causes of synchrony is important for predicting how spatial population synchrony and regional extinction probability will change with continued environmental change and habitat fragmentation. Recent studies indicate that global environmental change is affecting the frequency, intensity, spatial extent, duration, and timing of environmental patterns, ultimately changing the relationship between the environment and population dynamics (Di Cecco & Gouhier, 2018, IPCC, 2022). Most climate change scenarios predict a more synchronized climate in the future and a few studies have looked at the potential impact this climate change can have on spatial population synchrony (Post & Forchhammer, 2004, Defriez et al., 2016, Kahilainen et al., 2018). This will likely promote large-scale regional fluctuations in climate, which means we can also expect to see a concomitant increase in spatial population synchrony for species whose dynamics are highly environmentally driven (Post & Forchhammer, 2004, Nicolau et al., 2022). Increasing variability and severity of climatic events have been identified as the largest threat to population stability in birds (Møller et al., 2004) and insects (Harvey et al., 2023). Being able to predict species-specific responses to changes in environmental variability is an important tool in mitigating climate change impacts and avoiding population collapse. These sorts of generalizations shown in our results can aid managers to better make conservation prioritization decisions for species of conservation concern. Understanding these specific drivers of spatial population synchrony is important in the face of increasingly severe threats to biodiversity and could be key for successful future conservation outcomes.

## Supplemental Materials

**APPENDIX 1. Figure S1.**
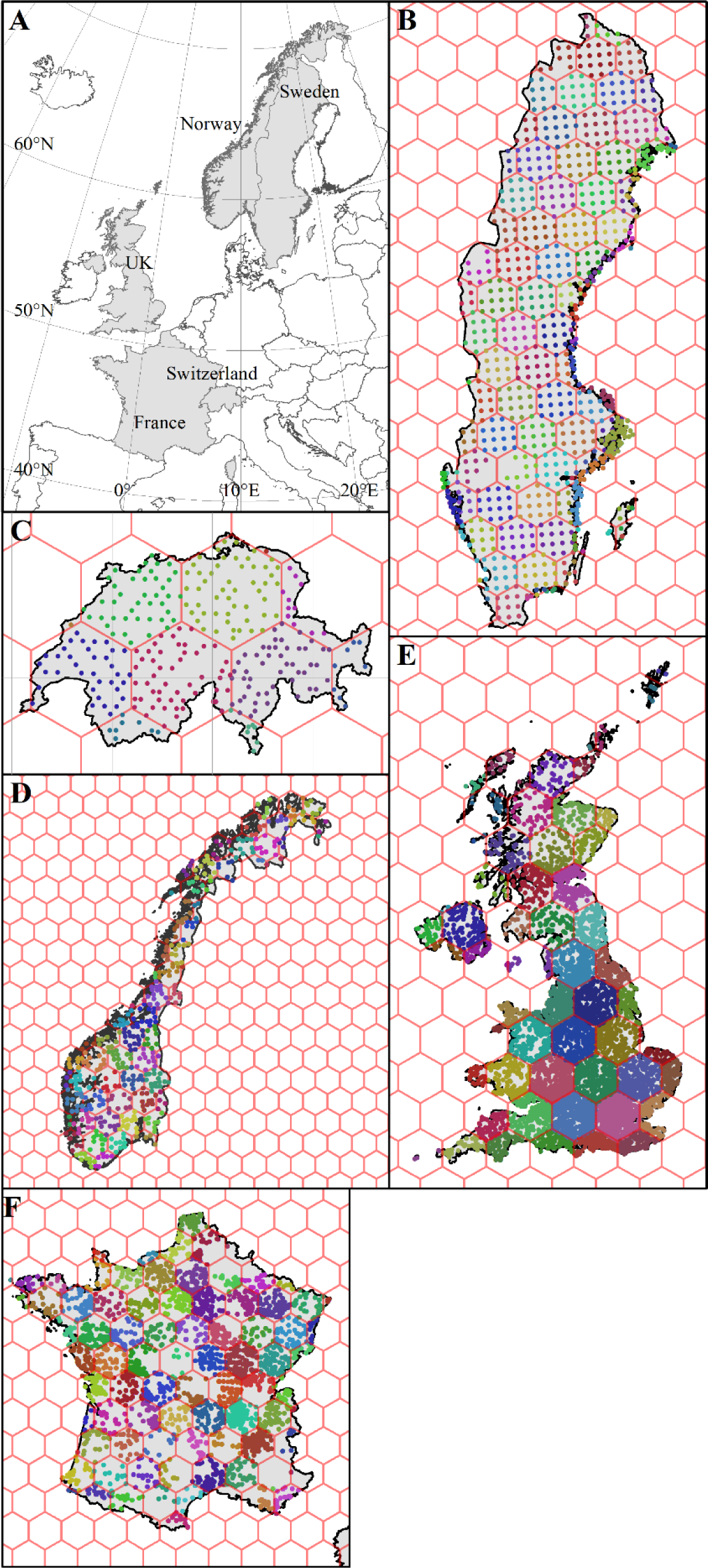
Figure of all study sites for birds in all countries. B) Sweden, C) Switzerland, D) Norway, E) the UK (United Kingdom), and F) France. 100km-diameter hexagonal grids used to aggregate survey sites shown in red.

**APPENDIX 1. Figure S2:**
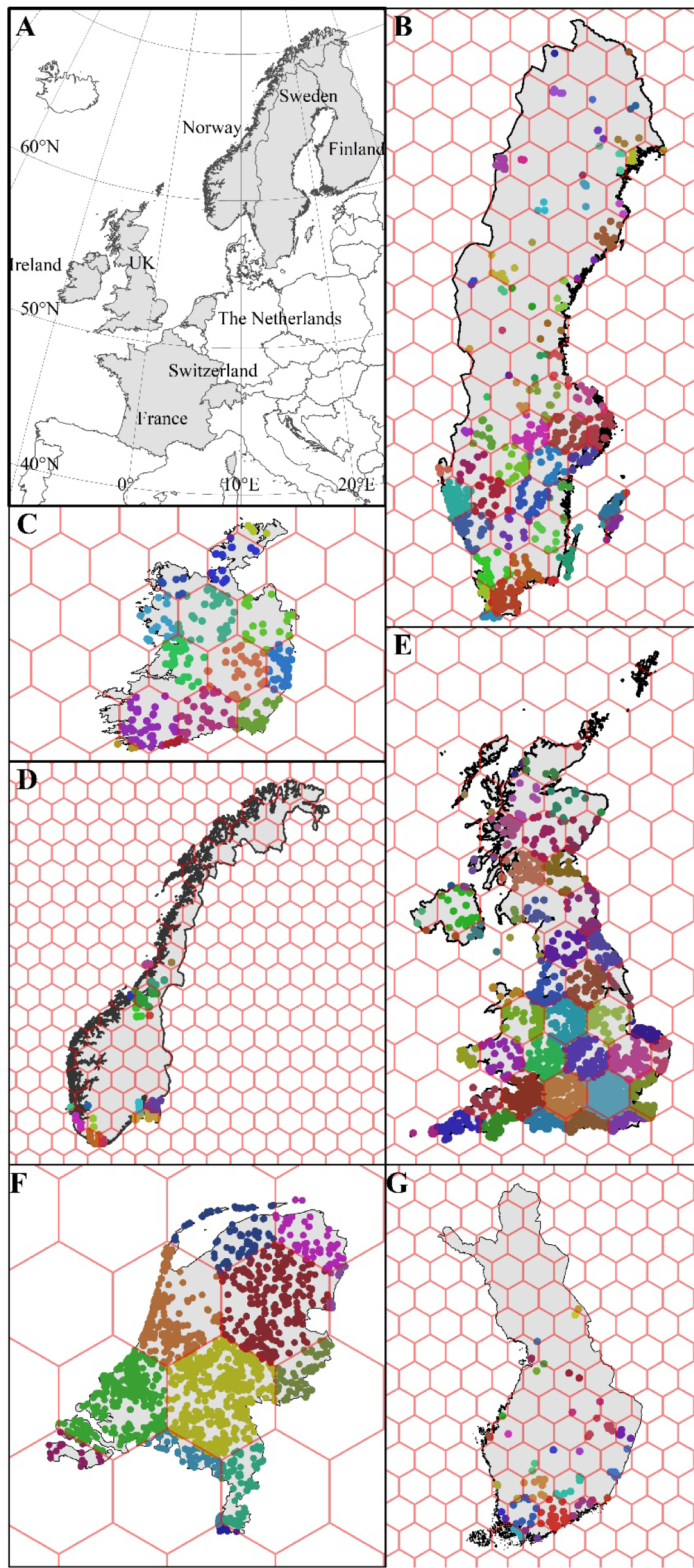
Figure of all study sites for insects in all countries. B) Sweden, C) Ireland, D) Norway, E) the UK (United Kingdom), F) the Netherlands, and G) Finland. 100km-diameter hexagonal grids used to aggregate survey points shown in red.

**APPENDIX 1. Figure S3.**
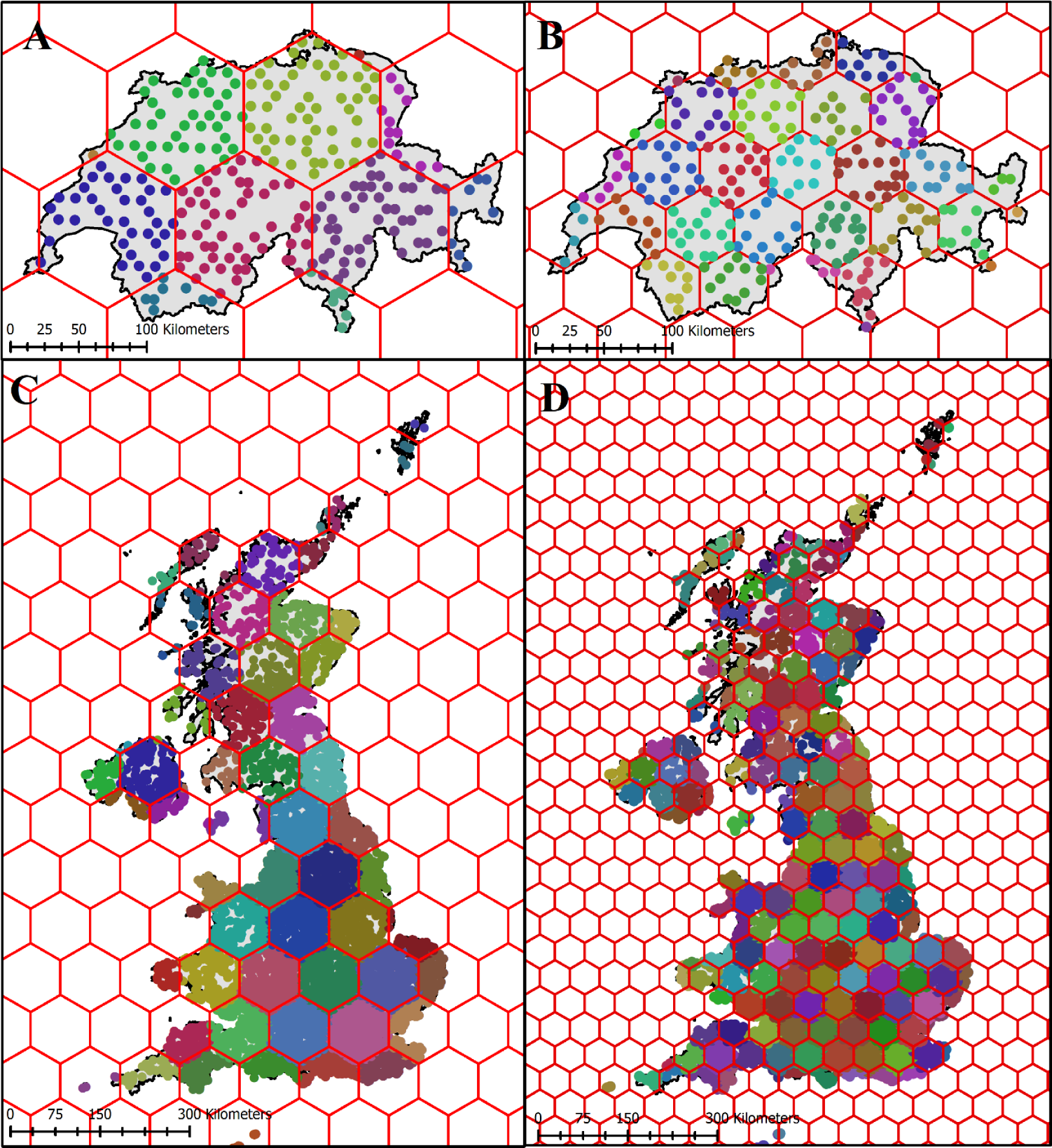
Switzerland (A, B) and the United Kingdom (C, D) grid size comparison. Panels A and C show point aggregation for an overlay grid of 100km diameter per grid cell. Panels B and D show point aggregation for an overlay grid of 50km diameter per grid cell.

**APPENDIX 2.**
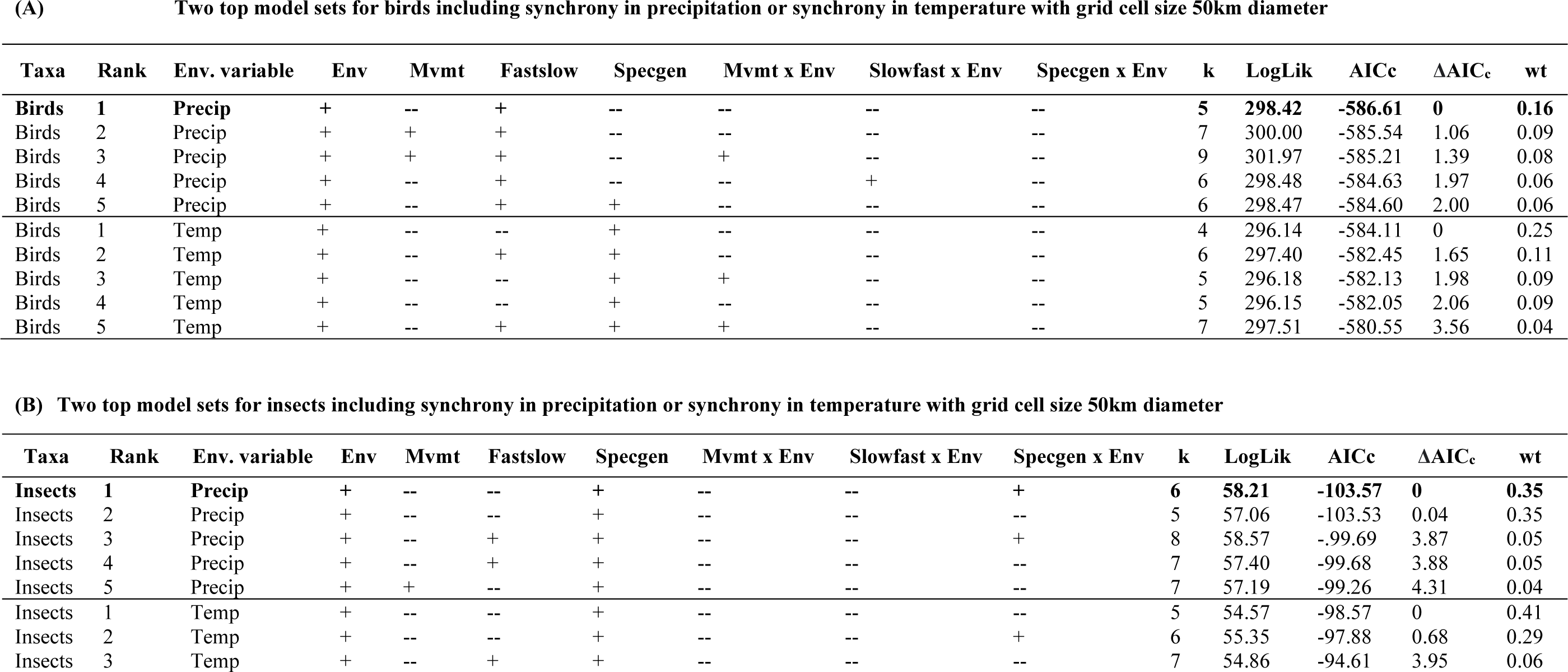
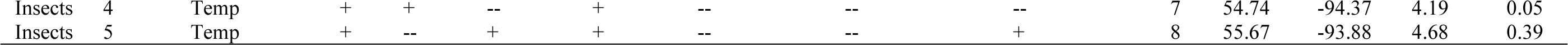
Model selection results for the analysis of spatial synchrony in annual population growth rates of (A) birds and (B) insects in Europe. Synchrony estimates are based on pairs of populations *≤*250km apart, merged in grid cells of size 50km diameter. Covariates designated with a “+” were present in model. Covariates included environmental synchrony (Env; in terms of mean summer precipitation [Precip] or mean summer temperature [Temp]), movement (Mvmt), fast-slow life history continuum (Fastslow), specialist/generalist (Specgen), and two-ways interactions between environmental synchrony and the life history traits. We also included a covariate for median distance at which synchrony was calculated (Med dist). Only one environmental covariate (precipitation or temperature) was included in each model because of collinearity, resulting in two different model sets. Only the top five models are presented (rank 1-5). We relied upon Akaike’s Information Criterion with a small sample size correction (*AICc)* for model selection and used Akaike model weights (*wt*) and *ΔAICc* to identify the top model. Number of parameters in model indicated by column k. LogLik = log-likelihood.

**APPENDIX 3.**
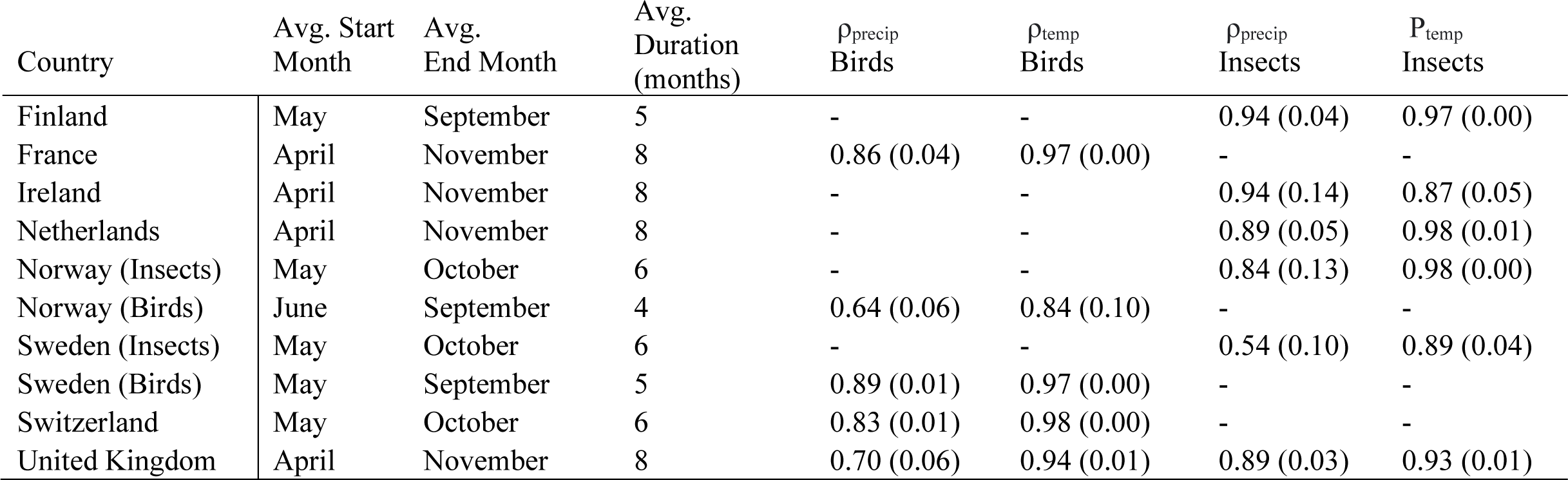
Country-specific mean (one standard error in parenthesis) within 250km of synchrony in precipitation (*ρprecip*) and temperature (*ρtemp*) with description of the periods included in the estimation. Months included had average temperature ≥ 5 degrees Celsius for the duration of the survey period.

**APPENDIX 4:**
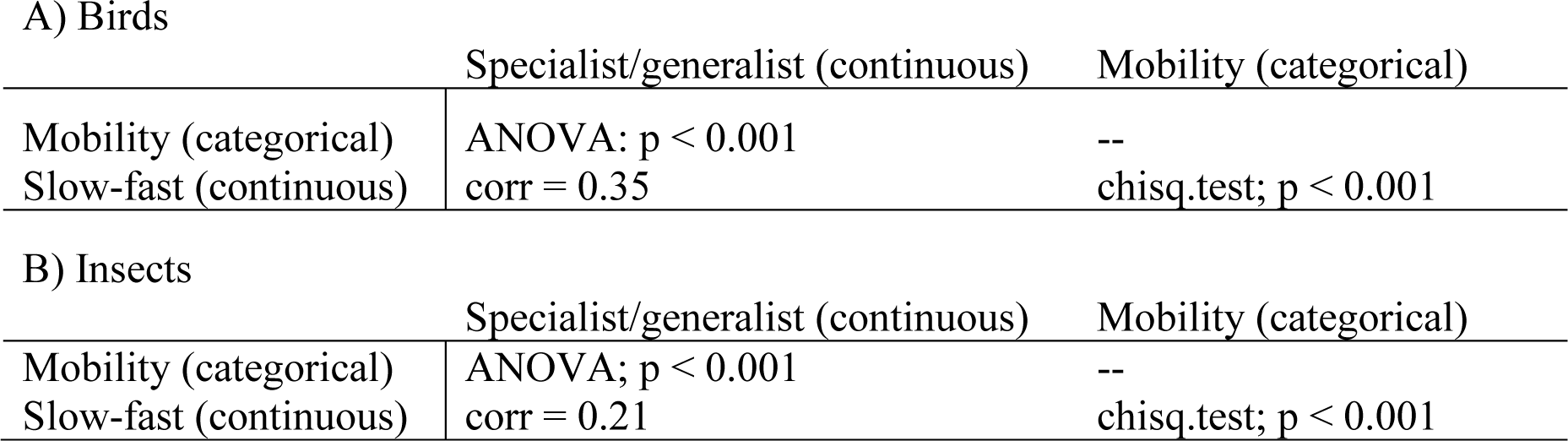
Correlations and dependencies between covariates used in analysis. For birds, position on the fast-slow life history continuum and mobility were not independent (chisq.test; p < 0.001). Position on the fast-slow life history continuum had a 0.35 correlation with specialist/generalist classification, while mobility and specialist/generalist classification were independent (ANOVA; p < 0.001). For insects, position on the fast-slow life history continuum was not independent from mobility (chisq.test; p < 0.001) and had a 0.21 correlation with specialist/generalist classification. Mobility and specialist/generalist classification were independent (ANOVA; p < 0.001).

**APPENDIX 5.**
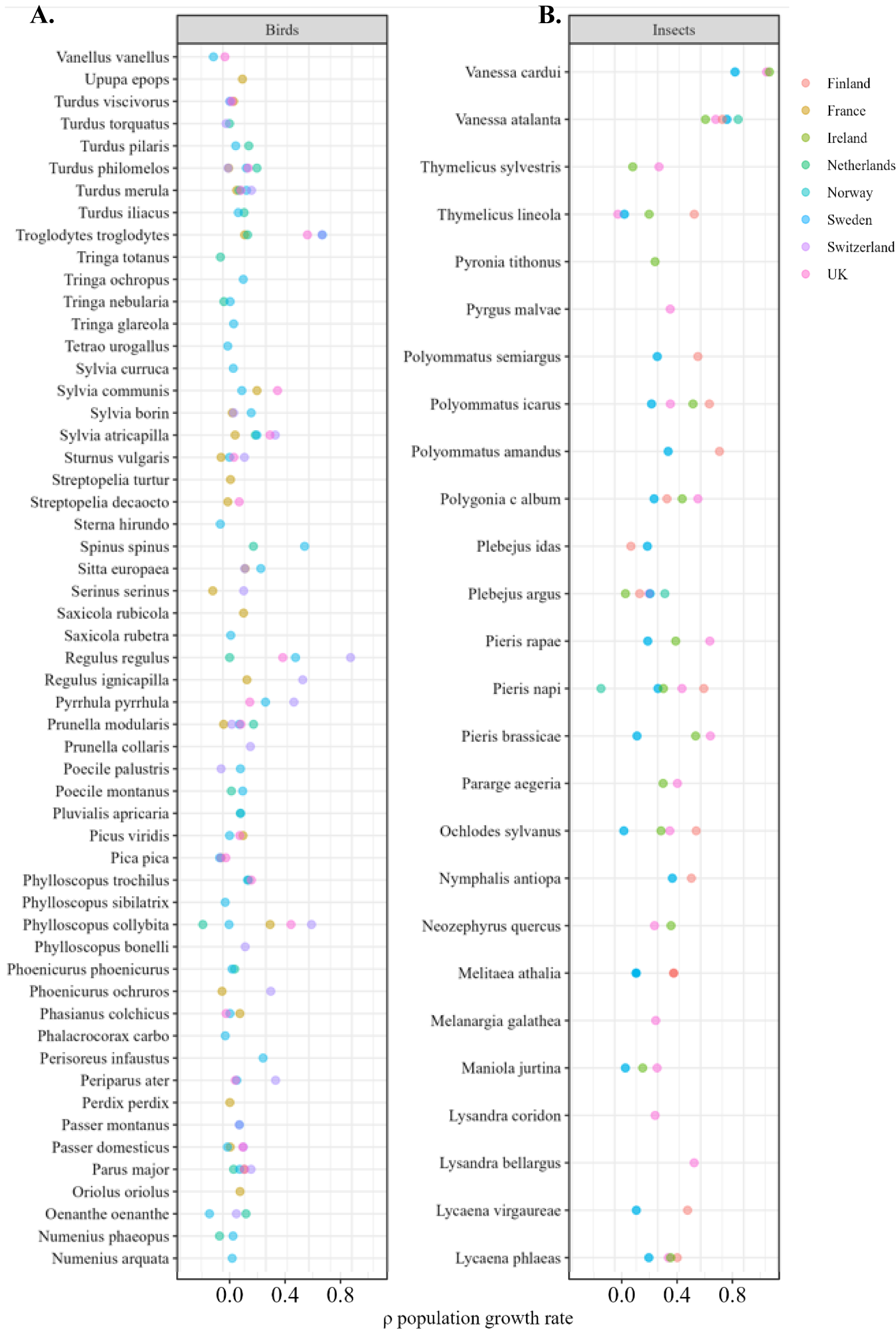
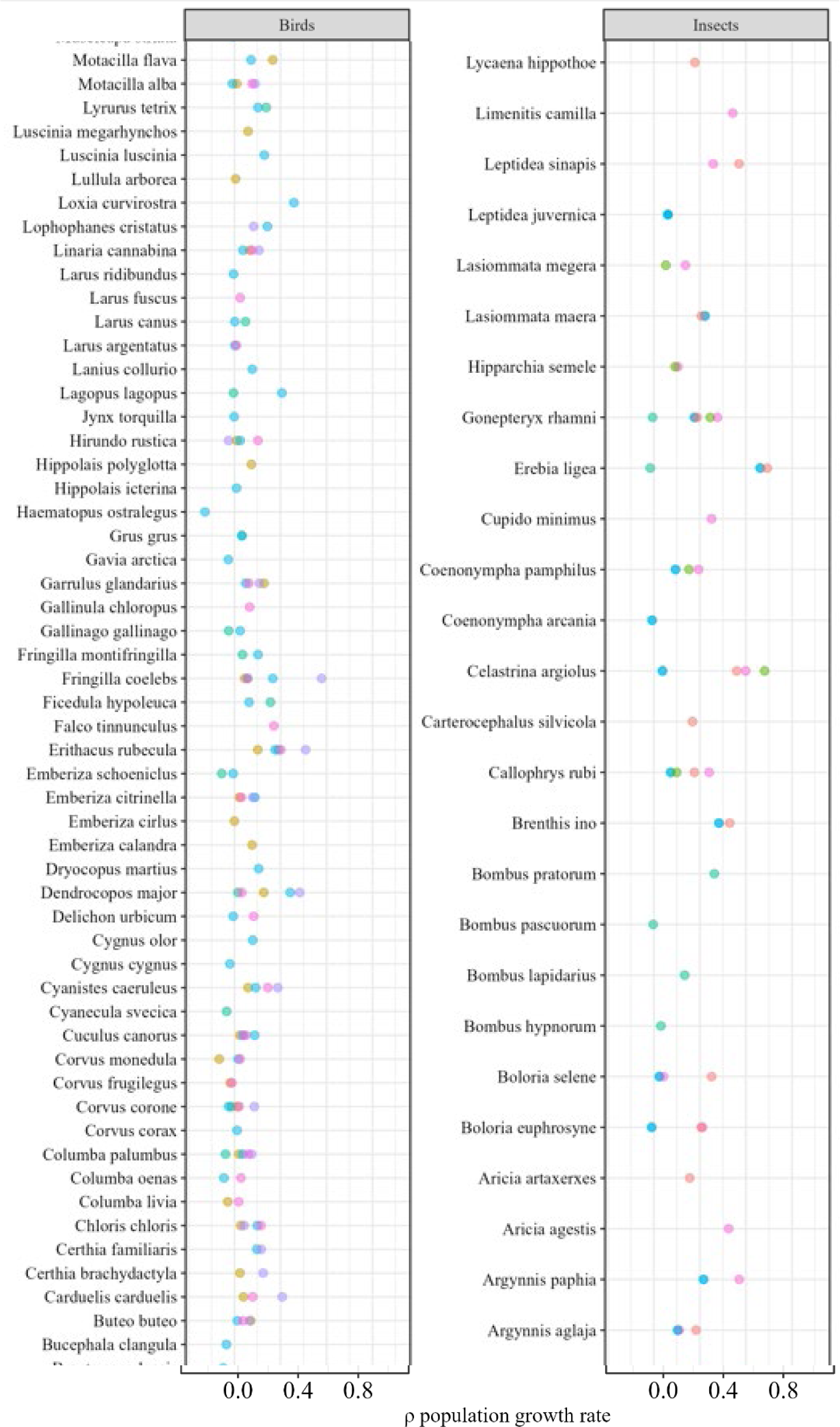
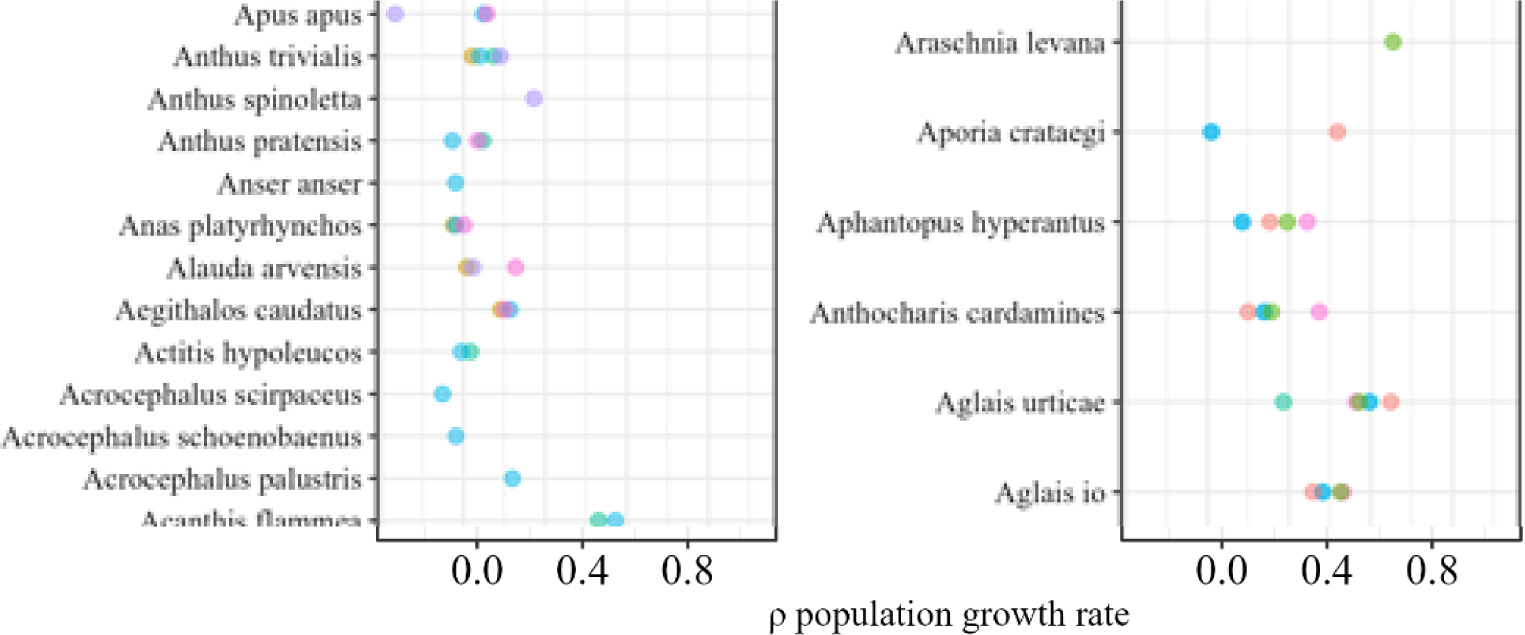
The distribution of country-specific mean spatial synchrony for different species of birds and insects. Mean synchrony in population growth rate was calculated as the average of the pairwise synchrony estimates between all pairs of grid cells within 250km of one another. Species that had three or more estimates of synchrony have distributions shown in grey. Country from which estimate comes indicated by point color. Distributions of synchrony are calculated from the R package *ggridges* function *geom_density_ridges*, which computes a kernel density estimate from the data.

**APPENDIX 6:**
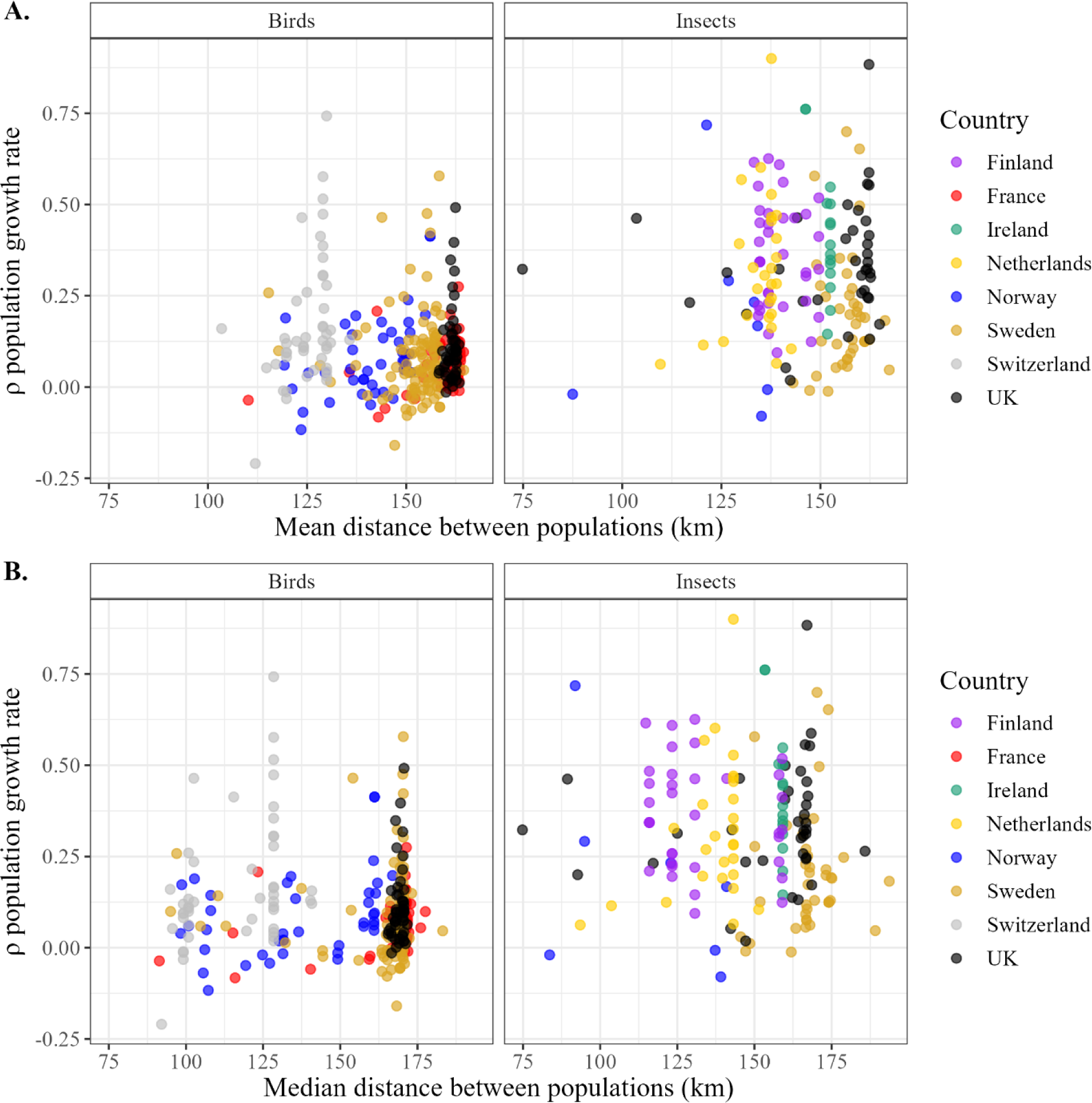
Mean (A) and median (B) distances (km) between populations used to calculate syncrony in population growth rates for all populations within 250km. Colored dots represent individual species’ mean distances separated by country and taxa.

